# New insights of the local immune response against both fertile and infertile hydatid cysts

**DOI:** 10.1101/390989

**Authors:** Christian Hidalgo, Caroll Stoore, Karen Strull, Carmen Franco, Felipe Corrêa, Mauricio Jiménez, Marcela Hernández, Karina Lorenzatto, Henrique B. Ferreira, Norbel Galanti, Rodolfo Paredes

**Affiliations:** Escuela de Medicina Veterinaria, Facultad de Ciencias de la Vida, Universidad Andres Bello, Santiago, Chile; Staff pathologist, Clinica Santa Maria, Santiago, Chile; Laboratorio de Biología Periodontal, Facultad de Odontología Universidad de Chile, Santiago, Chile; Facultad de Ciencias de la Salud, Universidad Autónoma de Chile, Santiago, Chile; Laboratório de Genômica Estrutural e Funcional, and Laboratório de Biologia Molecular de Cestódeos, Centro de Biotecnologia, UFRGS, Porto Alegre, RS, Brazil; Programa de Biología Celular y Molecular, Instituto de Ciencias Biomédicas, Facultad de Medicina, Universidad de Chile, Santiago, Chile

**Author notes:** **Corresponding author:** (RP).

## Abstract

**Background:** Cystic echinococcosis is caused by the metacestode of the zoonotic flatworm *Echinococcus granulosus*. Within the viscera of the intermediate host, the metacestode grows as a unilocular cyst known as hydatid cyst. This cyst is comprised of two layers of parasite origin: germinal and laminated layers, and one of host origin: the adventitial layer, that encapsulates the parasite. This adventitial layer is composed of collagen fibers, epithelioid cells, eosinophils and lymphocytes. To establish itself inside the host, the germinal layer produces the laminated layer, and to continue its life cycle, generates protoscoleces. Some cysts are unable to produce protoscoleces, and are defined as infertile cysts. The molecular mechanisms involved in cyst fertility are not clear, however, the host immune response could play a crucial role.

**Methodology/Principal fidings:** We collected hydatid cysts from both liver and lungs of slaughtered cattle, and histological sections of fertile, infertile and small hydatid cysts were stained with haematoxylin-eosin. A common feature observed in infertile cysts was the disorganization of the laminated layer by the infiltration of host immune cells. These infiltrating cells eventually destroy parts of laminated layer. Immunohistochemical analysis of both parasite and host antigens, identify these cells as cattle macrophages and are present inside the cysts associated to germinal layer.

**Conclusions/Significance:** This is the first report that indicates to cell from immune system present in adventitial layer of infertile bovine hydatid cysts could disrupt the laminated layer, infiltrating and probably causing the infertility of cyst.

**Author Summary:** Cystic echinococcosis is caused by the zoonotic flatworm *Echinococcus granulosus*. Within the viscera of the intermediate host, mainly liver and lungs of herbivores such as cows and sheep as well as human beings, the parasite grows as a unilocular cyst known as hydatid cyst. These cysts develop in their inner chamber a structure known as protoscolex, when consumed by the definitive host (e.g. dogs), it grows into a worm that resides in the small intestine and produces eggs that contaminate the environment. In cattle, most hydatid cysts are unable to produce protoscoleces, and thus are termed infertile hydatid cysts. The molecular mechanisms that explain the causes of hydatid cyst infertility remain unknown. We routinely collected cattle hydatid cysts from both liver and lugs and processed them for histological analysis. We found that there is a subset of fertile hydatid cysts that have low protoscolex viability and high immune infiltration surrounding the cyst. All infertile cysts have high immune infiltration, and many of them show disruption of the laminated layer and immune cells of host origin inside the cyst. This is the first report that shows that the cyst can be infiltrated by the host immune system.

## Introduction

Cystic echinococcosis (CE) is a major zoonotic disease caused by infection with the metacestode stage (hydatid cyst) of the flatworm *Echinococcus granulosus*. It has a worldwide distribution with an estimated 4 million people infected and another 40 million at risk [1]. High parasite prevalence is found in Eurasia, Africa, Australia and South America. CE affects more severely South American countries characterized by extensive grazing livestock farming including Argentina, Brazil, Chile, Peru, and Uruguay [2]. The life cycle of this parasite includes two mammal hosts. The definitive hosts are dogs and other canids; while ungulates and other mammals act as intermediate hosts [3] such as sheep, goats, cattle, pigs, buffaloes, horses and camels [4]. The most common infection targets are the liver and lungs because they are highly irrigated. Within these viscera, a unilocular cyst forms, that will grow gradually from 1 cm to 5 cm a year [5]. The hydatid cyst is circumscribed by a layer generated by the intermediate host in response to the parasite, named adventitial layer, which mainly consists of epithelial cells and connective tissue [6]. The adventitial layer can have variable thickness and may present some focal fibrosis as result of host immune response that considers the cyst a foreign body [7]. The lumen of the hydatid cyst is filled with the so called hydatid fluid and is surrounded by two layers of parasite tissue; the innermost layer is called germinal layer is cellular and is intimately attached to an acellular layer called laminated layer, the latter is in close contact with the adventitial layer. The germinal layer is composed by embryonic cells whose function is to elaborate the different elements of hydatid cysts [7]. These embryonic cells differentiate into buds that finally generate protoscoleces (PSC), the infectious parasite form for the definite host [5]. The laminated layer is generated by the germinal layer and is described as a specialized extracellular matrix that is found only in the genus *Echinococcus*. Macroscopically, it is seen as a whitish membrane, formed by various layers of mucopolysaccharides and keratin, evolutionarily adapted to maintain the physical integrity of metacestodes and to protect the cells of the germinal layer from host immunity [8]. In the intermediate host, it is possible to find two different types of hydatid cysts: fertile cysts, in which PSC are attached to the germinal layer and free into the hydatid fluid; and infertile cysts, without PSC. Infertile cysts are unable to continue with the parasite life cycle. The reason behind why two types of cysts are present remains unclear [9]. In many geographical areas, including Chile [10], cattle has been associated with low fertile hydatid cysts counts (<30%) in both *Echinococcus granulosus sensu lato* [11–14] and *Echinococcus granulosus sensu stricto* [15], so it is a suitable model to study cyst infertility mechanisms. Our research team, so far has been working in understanding the causes of infertile hydatid cyst in cattle, identifying both higher apoptosis levels in germinal layer of infertile cysts [9] and different immunoglobulin profiles [16]. Possible relations of the laminated and adventitial layers with fertility or infertility, however, have never been addressed. In this work, we present a systematic comparative study of both the laminated and adventitial layer in fertile and infertile hydatid cysts obtained from naturally infected cattle. Their morphohistological characteristics were described and compared, demonstrating the infiltration of host immune cells and providing evidences of their effect on and contribution to cyst integrity and fertility.

## Materials and Methods

### Sample collection, classification and genotyping

All bovine hydatid cyst samples were obtained at an abattoir in Santiago, Chile, as part of the normal work of the abattoir and with consent from both the veterinarians and the owners of the abattoir for sample collection. Both lung and liver samples were manually inspected by the official health office veterinarian and afterwards by our research team. This protocol study was approved by the Universidad Andres Bello Bioethics Board (protocol number 016/2016). For each positive viscera, the hydatid cysts were removed and placed in a sealed plastic bag and were stored at 4°C.

In the laboratory, each hydatid cyst was assigned a number, the hydatid fluid was aseptically aspirated and the cyst was opened along the longer axial plane. A sample of the germinal layer was observed in a conventional light microscope to determine if the cyst was either fertile or infertile. PSC viability was determined with trypan blue staining, and samples with viability lower than 90% were classified as low viability. Since smaller cysts (<1 cm in diameter) are still developing and have not started producing PSC, they were classified as “small cysts”. After the cyst fertility assessment, a sample of the cyst wall containing all three layers was fixated in Glyo-Fixx™ and embedded in paraffin. All samples were processed within 24 h of their procurement.

All hydatid cyst samples were genotyped using the method described by Bowles et al [17] in combination with sequencing of PCR products. Only samples from *Echinococcus granulosus sensu stricto* were used in this study.

### Haematoxylin-eosin morphohistological analysis

Paraffin blocks were cut to obtain 5 μm thickness sections and were stained with haematoxylin-eosin (H&E). Using an Olympus FSX100 microscope, each slide was examined to confirm that the three cyst layers were present. All infertile and small cyst samples that lacked the laminated layer were excluded from the analysis, however, all fertile cysts were studied due to the small sample size.

A seasoned pathologist examined blindly each slide, describing the germinal, laminated and adventitial layer features. For the adventitial layer, a score index from 0 to 3 was assigned to each cell type and inflammatory features grading its severity.

The thickness of the laminated layer was evaluated with the FSX-BSW software package that is included with the FSX100 microscope. The laminated layer length was measured in 20 consecutive areas in μm, obtaining both an average and its standard deviation.

For statistical analysis, Data was recorded in Excel 2010 datasheet and analyzed using RStudio IDE version 1.0.136 with R version 3.3.3. Outliers were identified using ROUT method with Q = 1% and differences were calculated using a two-way ANOVA with Tuckey’s post hoc analysis using. Statistical significance was considered with a P-Value <0.05.

### Immunohistochemical (IHC) analysis

To differentiate between parasite and host cells, we used two antibodies: one that targets *Echinoccocus granulosus* aldolase and another that targets host macrophages (Invitrogen S100A9 Monoclonal Antibody [MAC387]). Briefly, paraffin blocks were cut to obtain 3 μm thickness sections, and after deparaffinization and rehydration, for *E. granulosus* aldolase, antigen retrieval was performed with Citrate Buffer (10 mM sodium citrate, 0,05% Tween-20, pH 6.2), the primary antibody was incubated overnight at 4°C at a dilution of 1:1000. For cattle macrophages, antigen retrieval was performed by incubating slides with a 0.05% trypsin solution at 37°C for 15 minutes, the primary antibody was incubated 1 hour at room temperature at a dilution of 1:200. HRP-Conjugated secondary antibody (Jackson ImmunoResearch) anti-Rabbit (*E. granulosus* aldolase) or anti-mouse (cattle macrophages) was incubated for 1 h at room temperature at a dilution of 1:1000. Finally, DAB-Plus Substrate Kit (Life Technologies) was used for detection and slides were counterstained with haematoxylin.

## Results

### Bovine hydatid cyst sample distribution

From 2,961 bovines examined, 558 were positive to *Echinococcus granulosus* infection. After the paraffin embedding process and microscopic examination, 144 samples were obtained with the three layers in succession; 9 fertile (4 in lungs, 5 in liver), 129 infertile (91 in lungs, 38 in liver) and 16 small cysts (10 in lungs, 6 in liver). Fertile cyst viability ranged from 1% to 99% of viable PSC.

### The laminated layer thickness varies according to cyst location and type

Gross hydatid cyst examination of the laminated layer between fertile and infertile cysts was verified with microscopic measurement; fertile cysts have almost a six-fold thicker laminated layer than infertile and small cysts, 217 μm (± 18,4 μm) vs 33 μm (± 5,4), this difference is statistically significant (p <0.05). Conversely, small cysts have the laminated layer thickness similar to infertile cysts. There are also differences between cyst location; whereas fertile and infertile cysts have overall thicker laminated layers when found in the lungs (36 μm in lungs vs 29 μm in livers), small cysts have thicker laminated layers when found in the liver (26 μm in lungs vs 49 μm in liver), these differences are not statistically significant (p > 0.05) (Fig 1). The complete laminated layer measurements are available as S1 Table.

**Fig 1.**
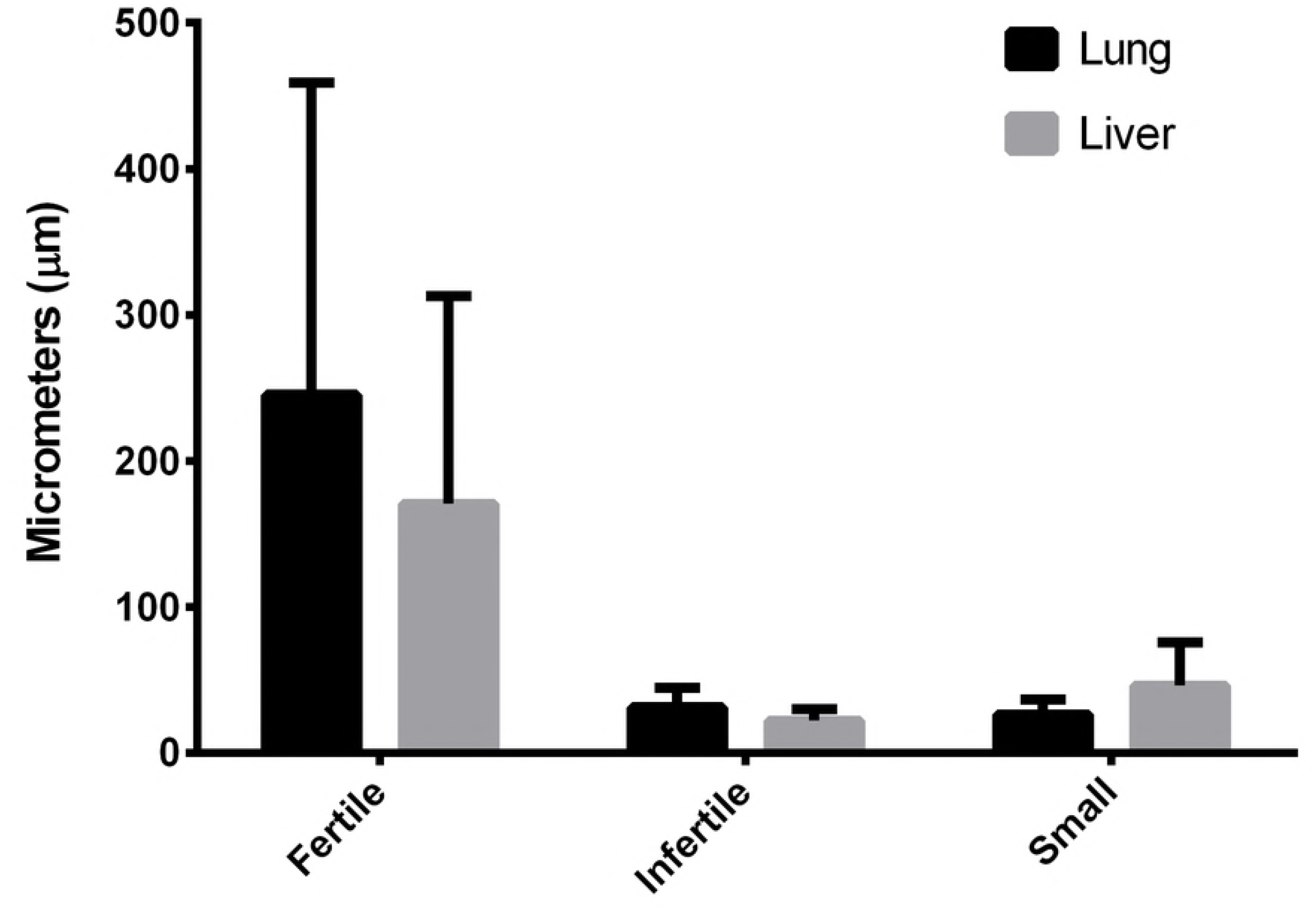
Laminated layer thickness (μm) according to hydatid cyst location and fertility. Data is shown as mean +/− standard deviation. Fertile, lung: n=4. Fertile, liver: n=5. Infertile, lung: n=83. Infertile, liver: n=36. Small, lung: n=10. Small, liver: n=6. The only statistical differences were between fertile and infertile/small cysts, regardless of organ localization.

### Fertile, infertile and small hydatid cysts present hallmark histological features in the adventitial layer

Fertile hydatid cysts with high PSC viability percentage, have germinal layer with PSC attached, cells in different developmental stages and thick laminated layers (>100 μm) that easily detach from the adventitial layer (Fig 2A); this adventitial layer is composed mainly of collagen fibers and fibroblasts (Fig 2B). Inflammatory cells, when present, are found beneath the collagen and fibroblast layer. There is a subset of fertile cysts that have low viability percentages (<10%) or zero viable PSC. When analyzed, these cysts have thinner laminated layers (<100 μm), and while the collagen and fibroblast layer is present in the adventitial layer, the inflammatory cells are found beneath the laminated layer (Fig 2C and 2D). Conversely, infertile hydatid cysts all share common features: the laminated layer is very thin, sometimes thinner than 5 μm. Beneath the laminated layer, all infertile cysts have palisading foamy macrophages. Supporting these macrophages, there are lymphoid follicles and multinucleated giant cells throughout the adventitial layer, with little presence of collagen fibers or fibroblasts (Fig 2E and 2F). Likewise, small hydatid cysts share the same histological features of infertile cysts (Fig 2G and 2H). The inflammation score is summarized in Fig 3 and full score data is available as S2 table.

**Fig 2.**
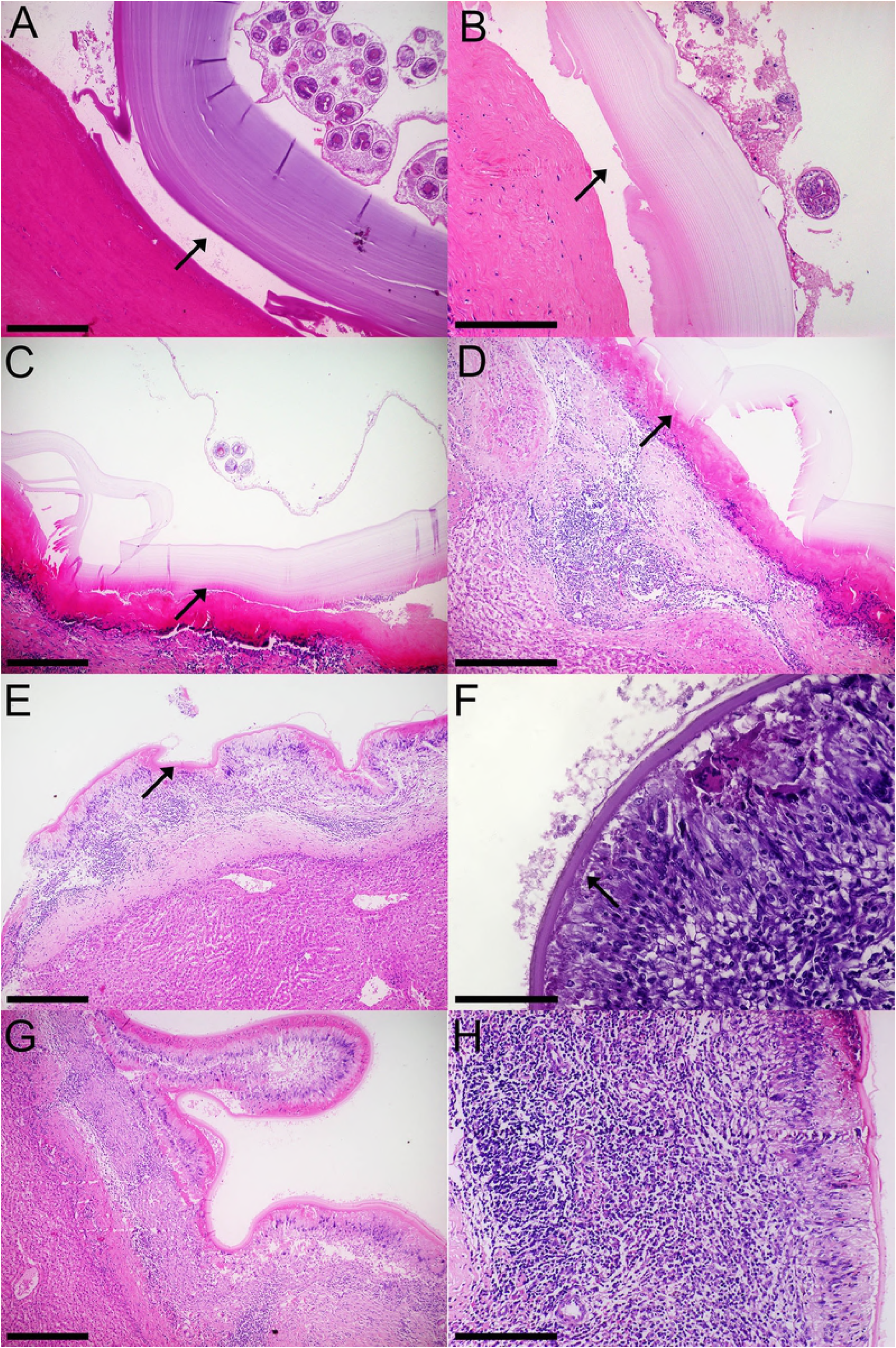
Histological characteristics of fertile, infertile and small bovine hydatid cysts from both lungs and livers. A, B: Fertile hydatid cyst with high viability protoscoleces (PSC) are characterized by a germinal layer with PSC, followed by a thick laminated layer and an adventitial layer composed of collagen fibers that detach from the laminated layer (arrows). C, D: Fertile hydatid cysts with low viability PSC feature the same germinal and laminated layer characteristic of high viability fertile hydatid cysts, but have an adventitial layer composed of inflammatory cells that are tightly attached to the laminated layer (arrows). E, F: Infertile hydatid cysts have little to none germinal layer cells with a thin laminated layer, the adventitial layer is composed palisading foamy macrophages, multinucleated giant cells and lymphoid follicles. This inflammatory infiltrate is tightly attached to the laminated layer (arrows). G, H: Small hydatid cysts (i.e. <1 cm) feature the same histological characteristics as infertile cysts. Size bar: A: 400 μm, B: 200 μm, C: 400 μm, D: 200 μm, E: 400 μm, F: 100 μm, G: 400 μm, H: 200 μm

**Fig 3.**
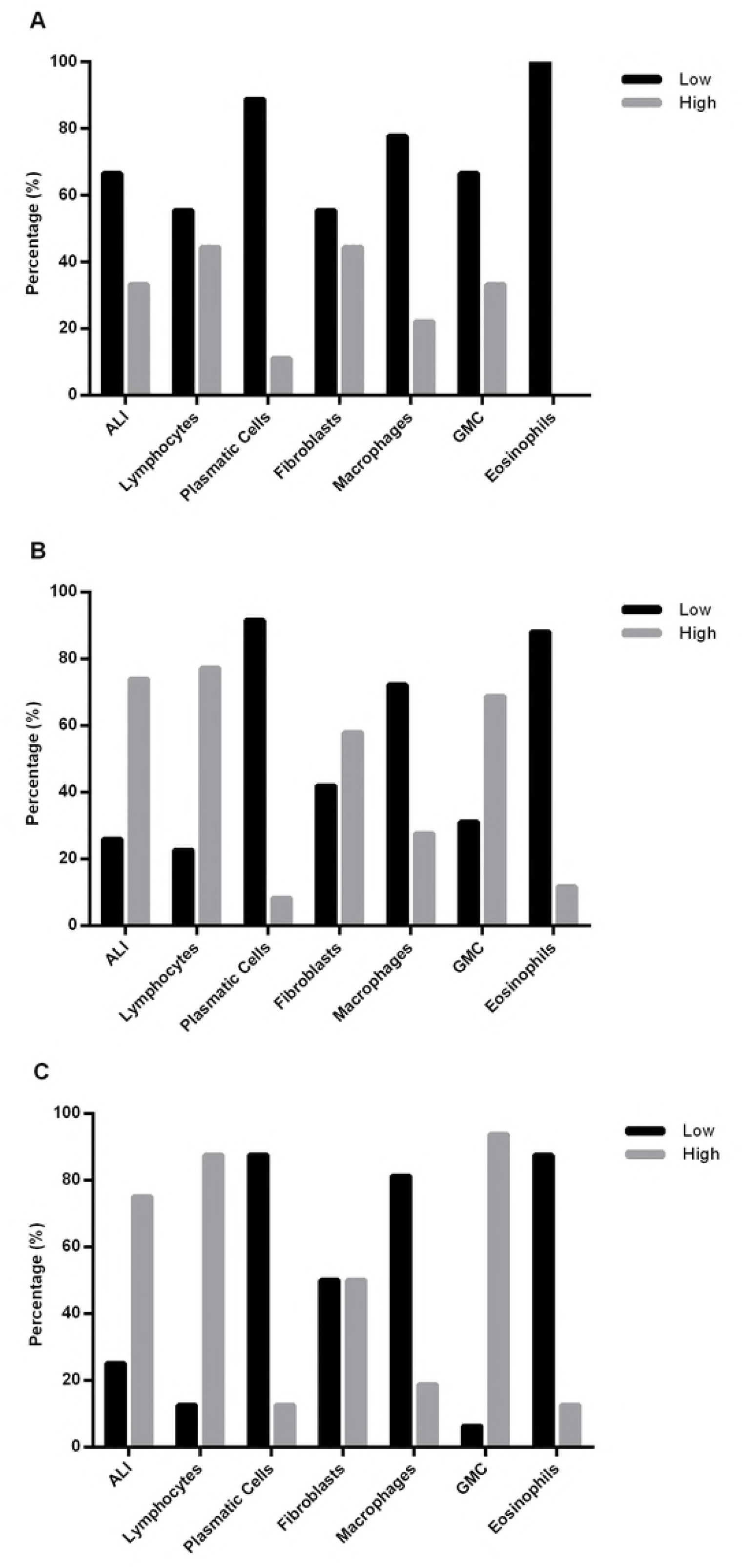
Inflammatory cell score between Fertile (A), Infertile (B) and Small (C) bovine hydatid cysts from both lungs and liver. Cell score ranged from 0 to 3 and was divided between low (0 to 1) and high (2 to 3) scores. Data is presented as a percentage of total samples. ALI = Adventitial layer inflammation; GMC = Multinucleated giant cells. Fertile n = 9, Infertile n = 129, Small n = 16.

A common feature, although not present in every infertile hydatid cyst, is the disorganization of the laminated layer by the infiltration of the host immune response; sometimes whole sections of the laminated layer can be found within the adventitial layer while in other samples the difference between laminated and adventitial layer becomes difficult to establish (Fig 4).

**Fig 4.**
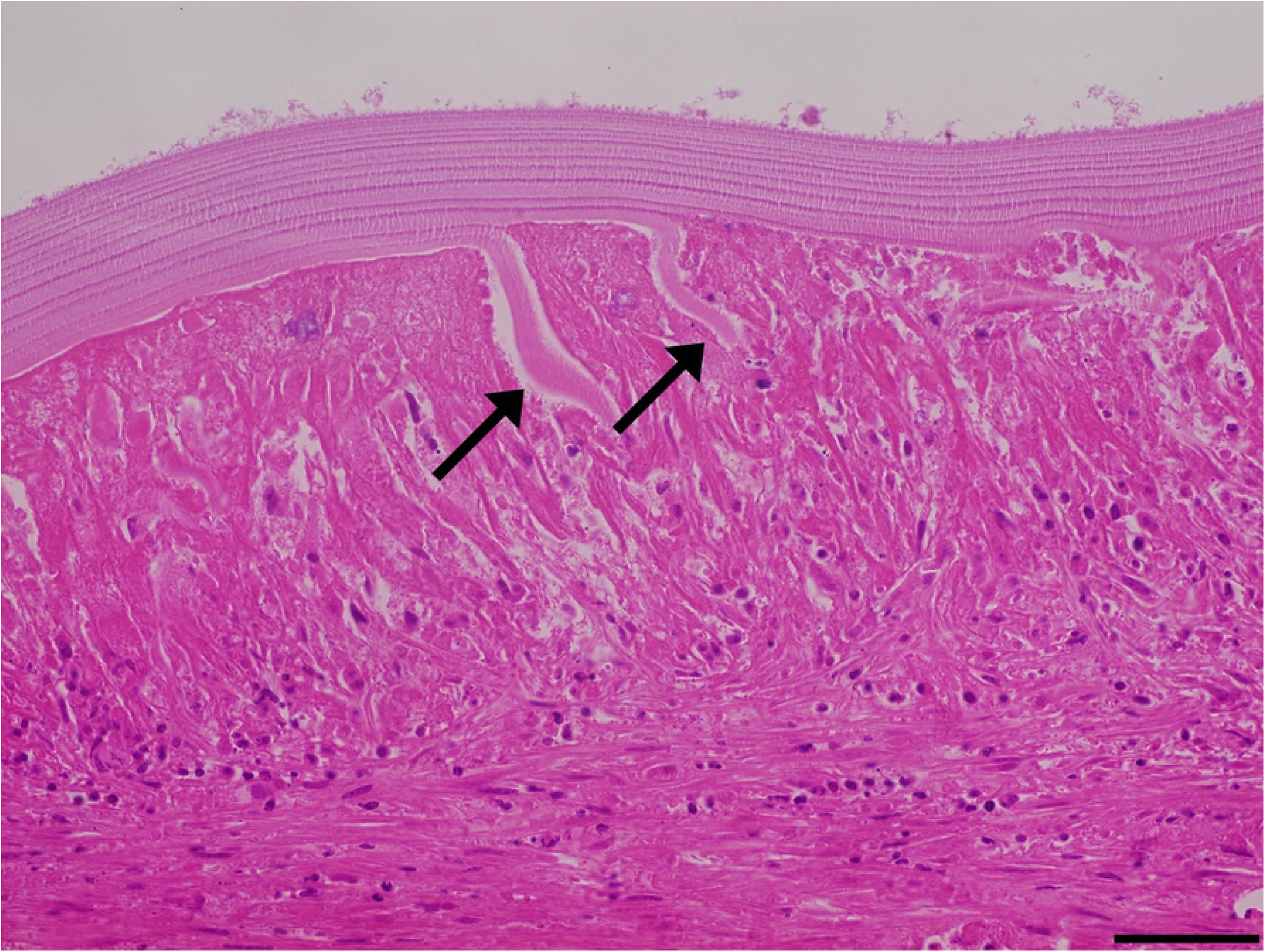
Laminated layer disorganization can be a common feature of low viability fertile cysts and both infertile and small hydatid cysts. The adventitial inflammatory cells infiltrate the laminated layer (arrows). Size bar: 50 μm.

### Infertile and small hydatid cysts present host immune cells inside the germinal layer

While analyzing the germinal layer of infertile cysts, there are many cells that have big nuclei, suggesting a mammalian rather than parasite origin. Sometimes these cells are present in big sheets or as single cells and resemble macrophages in the H&E analysis (Fig 5). To corroborate these results, IHC analysis of both host immune cells and parasite cells (Fig 6), demonstrates that these cells are not of parasite origin, as they are not detected by aldolase antibody (Fig 6C), but are positive for macrophage antibody (Fig 6F).

**Fig 5.**
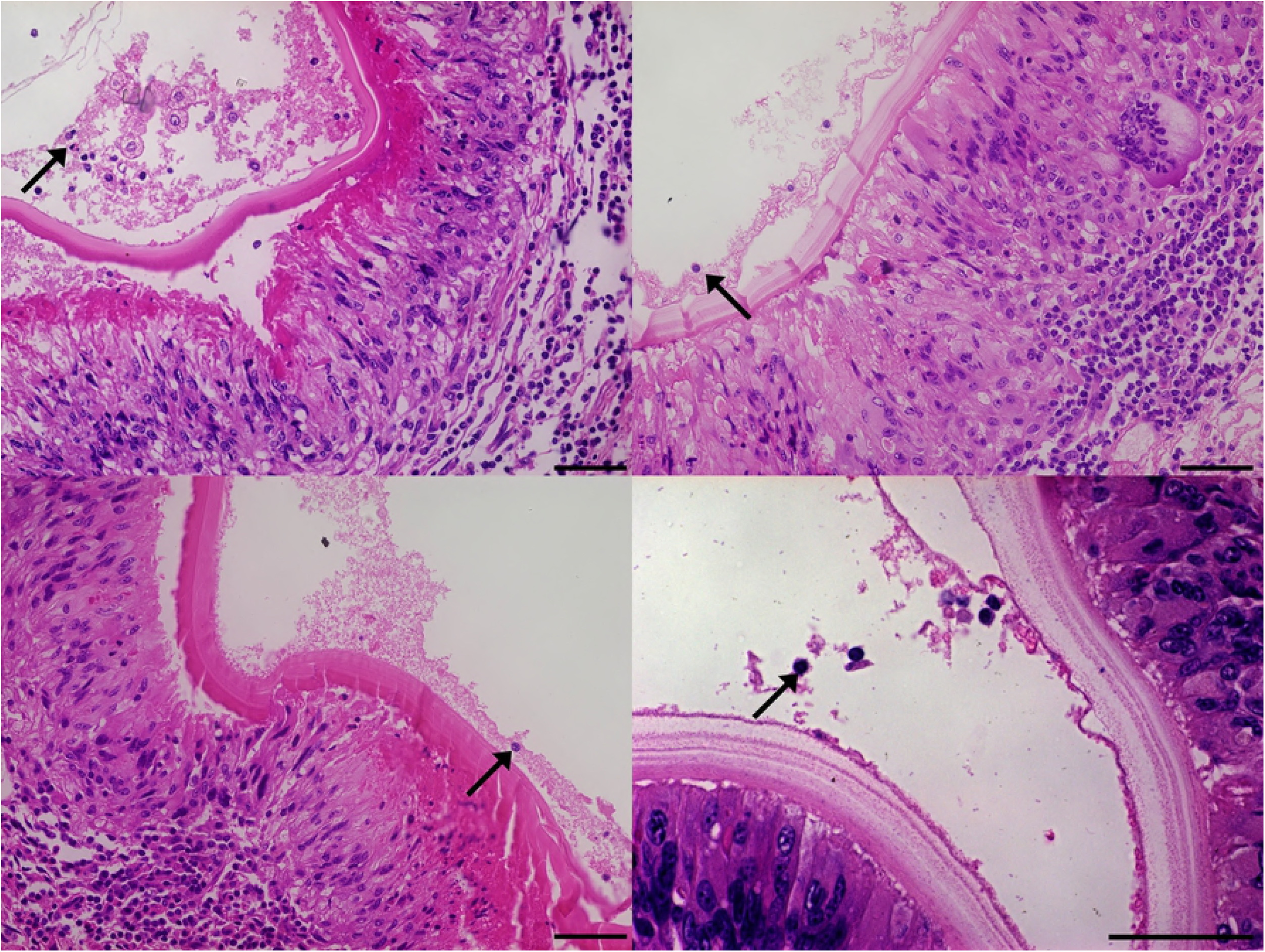
Infertile hydatid cysts have host cells infiltrating the germinal layer (arrows). Most of the infertile cysts without detachment of the laminated layer (LL) have the presence of host immune cells infiltrating the inner chamber of the cyst. Size bar 50 μm.

**Fig 6.**
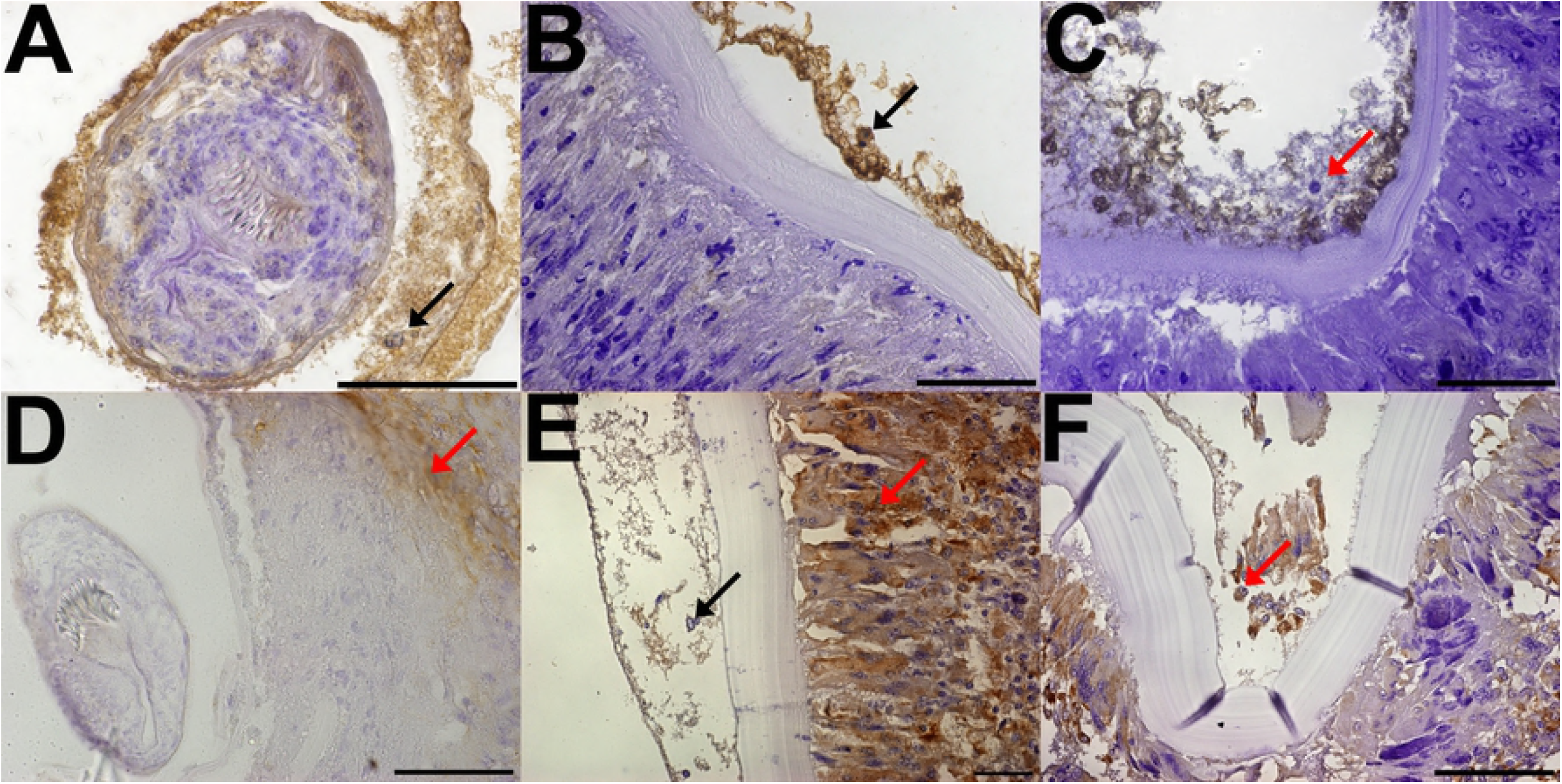
Immunohistochemical analysis of both fertile and infertile bovine hydatid cysts with *anti-Echinococcus granulosus* aldolase (A, B, C) and anti-macrophages antibodies (D, E, F). A: Fertile hydatid cyst, black arrow points at positive parasite cells inside the protoscolex. B: Infertile hydatid cyst without host cell infiltration, black arrow points at positive parasite cells along the germinal layer. C: Infertile hydatid cyst with host cell infiltration, red arrows points at host cells nucleus, which are negative to *Echinococcus granulosus* aldolase antibody detection. D Fertile hydatid cyst, red arrow points at positive macrophage while *Echinococcus granulosus* cells are negative E: Infertile hydatid cysts without host cell infiltration, red arrows points at positive macrophages while black arrow points at negative germinal layer cells. F: Infertile hydatid cyst with host cell infiltration, red arrow points at positive macrophages within both the germinal layer and the adventitial layer. Size bar: A, B, C, D, E: 50 μm. F: 100 μm.

## Discussion

*Echinococcus granulosus* infection elicits a granulomatous tissue reaction; characterized by the accumulation of cells of monocytic origin and are thought to be directed both at walling off and at eliminating the persistent foreign body. The hallmarks of granulomatous reactions are special types of activated macrophages called epithelioid cells and multinucleated giant cells [18].

The adventitial layer is usually described as a fibrous layer due to the host’s reaction to the parasite [6, 19–25], and several studies have described the cell composition of this layer. A study with fertile hydatid cysts found in the liver, showed that the adventitial layer has a significant amount of B lymphocytes, occasional polymorphonuclear cells and monocytes, however, they did no correlate this data with PSC viability, as the study was done in formalin-fixed, paraffin-embedded tissue samples [19]. Another study compared the differences in the adventitial layer between ovine and macropod hydatid cysts, with the adventitial layer of fertile cysts consisting of palisading macrophages with foamy cytoplasm and multinucleated giant cells or with granulation-type fibrous tissue devoid of a discernable covering epithelial layer [21]; and although the authors ponder whether cysts will develop and become fertile under inflammatory conditions, they do not correlate PSC viability with the characteristics they describe. The most complete study of the adventitial layer of bovine hydatid cysts was done by Sakamoto and Cabrera [6], where they describe that infertile cysts have lymphocytes, macrophages, granulocytes and polynuclear giant cells infiltrating the adventitial layer, with the smaller infertile cysts being surrounded by macrophage derived cells while eosinophils are involved in the response against larger cysts, and although they did exhaustive IHC analysis of the adventitial layer, surprisingly they did not report positive staining in the germinal layer of infertile cysts.

In this study, we analyzed both fertile and infertile hydatid cysts from liver and lungs of naturally infected bovine to identify characteristic histological features.

The thickness of the laminated layer was different between fertile and infertile hydatid cysts and also between liver and lungs tissue. It has been described that the parenchyma of these organs limits the growth rate of the hydatid cyst, with the lung being less dense that the liver [23], so it makes sense that the laminated layer is thicker in the lung cysts compared to liver cyst; small cysts on the other hand, had an inverse tendency; however, these differences were not statistically significant.

Although a fertile cyst is defined by the presence of PSC, we propose that a more complete definition should include viable PSC; as shown in this study, fertile cysts with low viability or dead PSC have adventitial layer characteristics of infertile cysts, and if sampled later, they possibly would be classified as such. However, because we work with natural infection samples, we are not able to confirm when the animals acquired the infection.

Many small hydatid cysts samples had adventitial layer characteristics similar to those of fertile cysts with dead PSC and infertile cysts. There is evidence that cysts from the same parasite strain, in the same organ and host can have differences in size, viability and fertility [25]. After examining 16 small hydatid cysts, all of which had adventitial layers with a strong immune reaction, we propose that small hydatid cysts with adventitial layer featuring palisading foamy macrophages, lymphocytes arranged in follicles, multinucleated giant cells, thin (<50 μm) laminated layer and host immune cells inside the germinal layer, should be regarded as infertile; while small cysts without these characteristic histological features could develop into either fertile or infertile cysts.

The laminated layer, which is secreted by the parasite contains mucins with O-type glycosylations and inositol hexakisphosphate (InsP6); these features are related to the parasite survival inside large mammals, and it has been proposed that the laminated layer is involved in down-regulating the local inflammatory response [26]. Also, it has been shown in mice that both macrophages and dendritic cells are activated by portions of the laminated layer [27]. In our results, many infertile and small hydatid cysts samples have clear signs of the host immune cells infiltrating the laminated layer and disorganizing it; with the eosin staining being more intense were this is happening; this could be due to the host macrophages secreting a combination of cathepsin K [18] and MMP-9 [28], although further experiments are needed to corroborate this.

Our results show that the bovine immune response, which is represented by the adventitial layer, is able in both lungs and liver, to disrupt the laminated layer of the hydatid cyst, and in many cases infiltrate the lumen, probably causing the destruction of the germinal layer promoting its infertility.

## Supporting Information Legends

S1 Table. Complete laminated layer measures (μm) of fertile, infertile and small bovine hydatid cysts, from both liver and lungs.

S2 Table. Inflammation scores (ranging from 0 to 3) of fertile, infertile and small bovine hydatid cysts, from both liver and lungs.

